# Differential roles of positive and negative supercoiling in organizing the *E. coli* genome

**DOI:** 10.1101/2022.12.30.522362

**Authors:** Ziqi Fu, Monica S. Guo, Weiqiang Zhou, Jie Xiao

**Affiliations:** Department of Biostatistics, Johns Hopkins Bloomberg School of Public Health, Baltimore, MD 21205, U.S.A.; Department of Microbiology, University of Washington, Seattle, WA 98198, U.S.A.; Department of Biophysics and Biophysical Chemistry, Johns Hopkins School of Medicine, Baltimore, MD 21205, U.S.A.

## Abstract

This study aims to explore whether and how positive and negative supercoiling contribute to the three-dimensional (3D) organization of the bacterial genome. We used recently published *Escherichia coli* GapR ChIP-seq and TopoI ChIP-seq (also called EcTopoI-seq) data, which marks positive and negative supercoiling sites, respectively, to study how positive and negative supercoiling correlates with the corresponding contact frequencies obtained from chromosome conformation capture sequencing (Hi-C and 5C). We found that supercoiled chromosomal loci have overall higher Hi-C contact frequencies than sites that are not supercoiled, with positive supercoiling surprisingly corresponding to higher spatial contacts than negative supercoiling. Additionally, Hi-C contact frequencies alone could identify positive, but not negative, supercoiling with high accuracy. The majority of positive and negative supercoils coincide with highly active transcription units, with a minor group likely associated with replication and other genomic processes. Our results suggest that both positive and negative supercoiling enhance chromosome interactions, but positive supercoils contribute more than negative supercoils to bring distant chromosomal loci closer in space. Based on these results, we propose new physical models of how the *E. coli* chromosome is organized differentially by positive and negative supercoils.

## INTRODUCTION

All living cells contain DNA that is compacted 1000-10, 000-fold in a nuclear space. In bacterial cells, DNA is packaged mainly through supercoiling, the twisting and crossing of the double-stranded DNA about itself, and the binding of nucleoid-associated proteins (NAPs), while eukaryotic cells use histones to organize their DNA within a nuclear envelope (1, 2). Supercoiling can be categorized as negative or positive, depending on the direction of the twists with respect to that of the DNA double helix, with both types of supercoiling forming coils (twists) or plectonemes (writhes) (3). Recently, chromosome conformation capture technologies such as Hi-C allow high-resolution interrogation of genome organization by measuring the physical interaction frequency of any two DNA fragments (4, 5, 6, 7). Despite revolutionizing our understanding of the overall architecture of chromosomes and revealing chromosome sub-structures such as DNA loops and topologically associated domains (TADs), Hi-C, like most other genomic technologies, captures DNA sequences but not DNA shapes. Consequently, how supercoils underlie the three-dimensional organization of chromosomes remains poorly understood.

In *E. coli*, previous work has revealed that the chromosome is organized into six ∼800 kilobase-long macrodomains, analogous to eukaryotic TADs, and that NAPs play differential roles in making short and long-range interactions (4, 5, 8). Our understanding of supercoiling in chromosome organization, however, is more limited. Supercoiling occurs during transcription as unwinding of the DNA leads to formation of positive supercoiling ahead of RNA polymerase and negative supercoiling behind it, referred to as the “twin-domain model” (9, 10). Replication also generates significant positive supercoiling ahead of the replisome (22). Recent experiments using psoralen, a DNA intercalator that specifically binds to undertwisted, negatively supercoiled DNA, indicates that negative supercoiling is pervasive in *E. coli* chromosomes (11) and organizes the local DNA (± 25 kb) around transcription units (12). These psoralen studies further suggest that the *E. coli* macrodomains may be organized by supercoiling. However, because psoralen only interrogates negatively supercoiled DNA, the contribution of positive supercoiling to these organizational effects can only be indirectly inferred.

Two recent supercoiling-mapping technologies, GapR-seq (13) and EcTopoI-seq (14), have largely overcome the technical limits of the previous psoralen approach, generating high-resolution (<1 kb) snapshots of the location and magnitude of positive and negative supercoiling, respectively. Both technologies rely on chromatin immunoprecipitation sequencing (ChIP-seq) to probe the genomic locations at which two shape-recognizing proteins, *Caulobacter crescentus* GapR and *E. coli* Topoisomerase I (EcTopoI) bind, respectively. GapR was shown to preferentially bind to overtwisted, positively supercoiled DNA and serve as a sensor for such DNA in *Caulobacter*, *E. coli* and yeast (13, 15). Similarly, EcTopoI, an enzyme that binds and removes excessive negative supercoils (16), was shown to localize to regions of negative supercoiling (14). GapR-seq and EcTopo-seq discriminate between these two DNA shapes and have an inverted enrichment pattern around transcription units, where EcTopo-seq labels negative supercoiling upstream of transcription while GapR-seq identifies positive supercoiling downstream of transcription, validating the twin-domain model and providing a whole-genome portrait of supercoiling (14).

In this study we computationally integrate GapR-seq, EcTopoI-seq, RNA-seq and Hi-C/5C datasets to describe how positive and negative supercoiling contributes to the three-dimensional organization of the *E. coli* genome (**Fig. 1A**). Our statistical analyses reveal that genomic regions containing supercoiling are associated with higher-frequency spatial interactions than those not containing supercoiling. These supercoiling-associated contacts can span entire macrodomains and have not been previously reported. Strikingly, we discovered that positive supercoiling is correlated with higher spatial contact than negative supercoiling. Using a machine-learning model, we found that positively, but not negatively, supercoiled loci can be accurately predicted from Hi-C data alone, suggesting that positive supercoiling is a major driver of chromosome architecture. Lastly, we observed that while most supercoiled loci colocalize with active transcription units as expected, a minor set does not, suggesting additionally genomic processes in generating supercoils and contributing to chromosome packaging. Our results suggest that positive supercoiling is pervasive and plays a dominant role in global genome organization.

**Figure 1.**
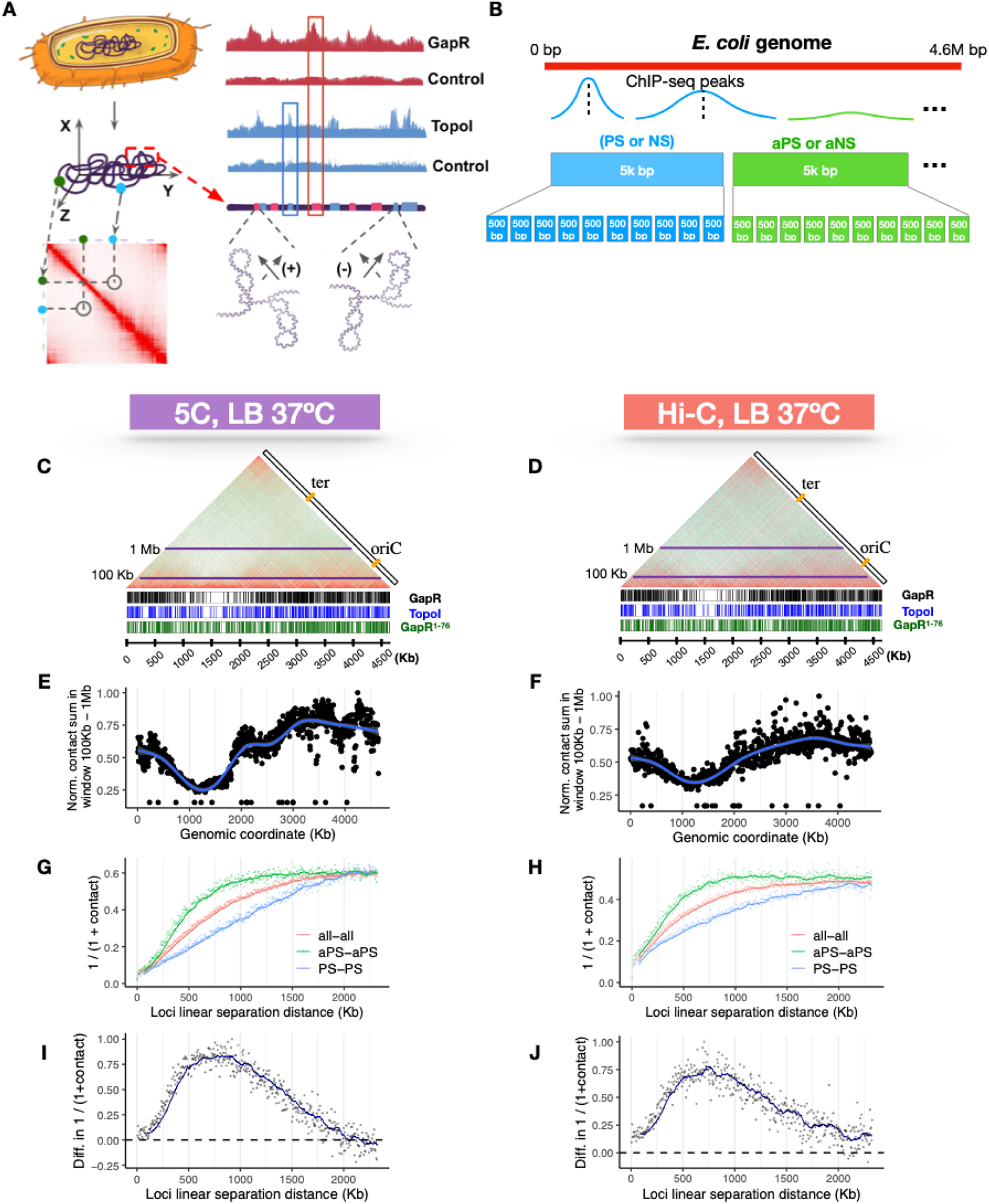
*E. coli* chromosomal loci containing positive supercoiling sites exhibit higher-than-average spatial contacts in Hi-C maps. **(A)** A cartoon illustration of the study design. We correlate the spatial contact frequency of any two *E. coli* chromosomal loci (green and cyan dots on the wiggle lines) as depicted in the Hi-C map (red square, bottom) with the supercoiling states of the two loci as identified by GapR (indicating positively supercoiled DNA) or Topo I (indicating negatively supercoiled DNA) chromatin-immunoprecipitation sequencing data (ChIP-seq, right). **(B)** A cartoon illustration of GapR/Topo I ChIP-seq data bin assignment. The 4.6 Mb *E. coli* genome is divided into 5 Kbp or 0.5 Kbp bins, with each bin assigned as PS (positive supercoiling site marked by GapR-peak), aPS (absent of a positive supercoiling site, no GapR peak), NS (negative supercoiling site marked by Topo I-peak), or aNS (absent of a negative supercoiling site, no TopoI peak), based on GapR-Seq and TopoI-seq data. A bin can contain either one, both, or none. **(C-D)** Comparison of genomic locations of GapR and TopoI peaks with respect to their spatial contacts with other genomic locations in the Hi-C contact heatmaps of *E. coli* MG 1655. Normalized (log2) contact map are plotted as symmetric halves from the 5C data set (LB, 37 °C, 5 Kbp resolution) generated by Lioy et. al (left) and Hi-C data set (LB, 37°C, 5 Kb resolution) generated in this study (right), with a 45% counterclockwise rotation. Vertical bars at the bottom of the contact maps mark the ChIP-seq peaks for GapR (black), GapR^1-76^ (green), and Topo I (blue). The two horizontal lines in purple (10Kb, 1Mb) represents the window used to compute the contact sum (in **E-F**) at each genomic locus in the contat map (5 Kb bin). The two reference rules provide the *E. coli* genomic coordinates (bottom, increment = 500 Kb) and locations of *Ori*C and *ter*C. **(E-F)** Normalized (log2) sum of the total contact counts in the 100 Kb and 1 Mb window of the contact map (C or D) at each locus (black dots) is plotted along the *E. coli* genomic coordinates. Blue curve shows the moving average (window size = 15) of the normalized contact sum and peaks near ∼3-4 Mb where *OriC* resides. **(G-H)** Inversed average contact frequencies of two genomic loci (*y*-axis) at each linear genomic separation distance (*x*-axis) with simple moving averages (colored curves, window size = 15) for loci pairs both containing positive supercoiling sites (PS-PS, blue), not containing positive supercoiling sites (aPS-aPS, green), and all loci pairs (all-all, red) show that PS-PS loci pairs have on average lowest inversed average contact frequencies (highest spatial contact) at all linear genomic distances than aPS-aPS or all-all loci pairs. **(I-J)** Numeric difference in the inversed average contact frequencies (y-axis) between PS-PS loci and aPS-aPS loci at each genomic separation distance (x-axis) reaches maximum in the range of 500-1000 Kbp.

## MATERIAL AND METHODS

### Hi-C/5C data processing

We align the Hi-C/5C sequence reads to the *E. coli* reference genome NC_000913.3 and follow the Hi-C-Pro 3.1.0 pipeline (17). The resolution is 5 Kb for the data set generated in this study and the one by Lioy et. al., and it is 0.5 Kb for the data set generated by Cockram et al. The contact count matrices are normalized using the iterative correction and eigenvector decomposition (ICED) method (18). The ICED contact counts are log2 transformed after adding a pseudo count. Then, we use the R package ComplexHeatmap (19) to produce all heatmaps.

### ChIP-seq data processing and analysis

For the two ChIP-seq data sets, we first perform a quality check to the sequence reads using FastQC (20) and eliminate adapters using Trimmomatics (21). Bowtie2 (22) is used to align all sequence reads to the *E. coli* reference genome NC_000913.3, follow by conversion of SAM to BAM files using Samtools (23). ChIP-seq peaks are detected using MACS2’s callpeak function (24). To obtain a set of reproducible ChIP-seq peaks, a customized R script (see **Data Availability**) is used to obtain peak signals across replicates.

### Hi-C/5C bin classification based on ChIP-seq signals

A 5 Kb genomic bin in the Hi-C/5C data is classified as PS (positive supercoiling site) or NS (negative supercoiling site) if it contains the center of at least one GapR or TopoI peak, respectively. The complementary set of genomic bins to the set of PS and NS bins are labeled as aPS (absent of a PS-site) and aNS (absent of a NS-site), respectively. Bins that contain only GapR peak centers but no Topo peak center are defined as PS-only (positive supercoiling only) bins, while NS-only (negative supercoiling only) bins contain only Topo peak centers but no GapR peak center. Bins that lack any peak center are labeled as aa (absent of either). Similarly, a genomic bin of size 0.5 Kb is classified as PS or NS if it has a non-empty overlap with at least one GapR or TopoI peak, respectively. Conversely, aPS and aNS are assigned to the set of bins that do not overlap with any GapR or TopoI peaks, respectively. This assignment strategy, which is more relaxed compared with the one used for the 5 Kb bins, is implemented to resolve severe class imbalance issues caused by false negative identification of PS and NS. **Supplementary Data** Table S1 and Figure S1D display the summary statistics for the number and the width of the peaks.

### Training the random forest models

We align Hi-C/5C sequence reads to the E. coli reference genome NC_000913.3, following the Hi-C-Pro 3.1.0 pipeline (27) with iterative correction and eigenvector decomposition (ICED) normalization. The resulting contact counts are log2-transformed with a pseudo count to form a contact frequency matrix. To prepare the feature matrix, we transform the contact frequency matrix such that its diagonal becomes the middle column, representing the self-contacts. Each column represents a genomic separation distance from a given locus (row), which serves as a feature for our model. The dimension of the feature matrix is 928 by 928, determined by the maximum Hi-C resolution. The response columns, or target vectors, are labeled according to each locus’ classification as PS/aPS and NS/aNS, as described in the bin classification step. Each row represents a genomic locus with a binning size of 5 Kb, and the model predicts whether this locus contains any GapR or TopoI peaks. It is worth noting that the genomic location of each bin (or row) remains inaccessible to the models, as we generated the testing sets and their corresponding training sets randomly. In other words, each row is considered as an independent data point by the models.

We then build separate models for predicting positive and negative supercoiling. To train these models, we partition the feature matrix into four balanced and non-overlapping sets and build four series of models using each of the quarters as testing data. For each testing set, we build random forest models by using 25%, 50%, or 75% data from its complement in the feature matrix as the training data. The process was repeated 10 times for each training data set. We use the R package caret (25) for random forest training, and each random forest is trained using 5-fold cross-validation with 10 repeats. We optimize the mTry parameter (from 2 to 42 by a step of 5) and nTrees parameter (from 200 to 1200 by a step of 100) using the receiver operating characteristic (ROC) metric.

### Evaluating random forest model prediction performance

After training each model, we evaluated its performance by making predictions on an independent testing set. That is, we ask the model to predict the PS/aPS and NS/aNS labels of an unseen set of loci using only their Hi-C frequencies. For each testing dataset and prediction percentage (25%, 50%, and 75%), we made predictions using the 10 trained models and computed the average AUROC and AUPRC. The AUPRC metric was used in addition to AUROC to account for any potential class imbalance issues (26).

AUROC (Area Under the Receiver Operating Characteristic Curve) is a created by plotting the true positive rate against the false positive rate for different threshold values. While AUROC may take any values between 0 to 1, AUROC = 1 indicates a perfect model and AUROC = 0.5 indicates a random classifier, which typically serves as a theoretical baseline performance. AUPRC (Area Under the Precision-Recall Curve) is created by plotting the ratio of true positive predictions to the total number of positive predictions (precision) against the ratio of true positive predictions to the total number of actual positive cases (recall). The AUPRC ranges between 0 and 1. AUPRC = 1 indicates a perfect classifier, while its baseline score is the ratio of positive cases. AUPRC = 0 indicates that the model’s precision is always zero, regardless of the recall. Both metrics measure the classification performance of the random forest model, and AUPRC is a more reliable metric in class imbalance cases.

To provide a baseline for comparison, we generated a set of models using the same feature matrices as the actual models, but with randomly permuted response/target vectors. These models were then tested on unseen testing sets that were correctly labeled (the same as the actual models). The training process for these baseline models was the same as that used for the actual models. We calculated the AUROC and AUPRC for these baseline models and use them as references.

### Growth conditions and chemical treatment for *E. coli* used in Hi-C

*E. coli* was grown in LB (10 g/L NaCl, 10 g/L tryptone, 5 g/L yeast extract) at 37 °C with shaking at 200 rpm. Optical density was measured at 600 nm using a Genesys 10 Bio Spectrophotometer.

### Chromosome conformation capture deep-sequencing (Hi-C)

Hi-C experiments were performed as described previously with the following modifications (7). Wild-type *E. coli* K-12 MG1655 was grown to OD600 ∼0.2 and then fixed with 3% formaldehyde for 10 min at room temperature before quenching with 0.375 M final concentration glycine for 15 min. Fixed cells were pelleted, washed twice with 1X M2 salts (6.1 mM Na_2_HPO_4_, 3.9 mM KH_2_PO_4_, 9.3 mM NH_4_Cl), then washed 2 more times with 1X TE buffer (10 mM Tris-HCl [pH 8.0] and 1 mM EDTA), before resuspension in 1X TE at ∼10^7^ cells / µL. Resuspended cells were divided into 25 µL aliquots before storage at -80 °C. Each Hi-C experiment was performed with 2 x 25 µL aliquots. To each 25 µL aliquot, 20 000 U of Ready-Lyse Lysozyme (Lucigen) was added, and the mixture was incubated for 30 min before addition of SDS to a final concentration of 0.25% for 15 additional minutes to completely dissolve cell membranes and release chromosomal DNA.

Each aliquot of DNA was then digested with HpaII (NEB) as follows. A 50 uL reaction was assembled containing lysed cells, rCutSmart buffer, and 5 µL of 10% Triton-X 100, which was allowed to sit for 15 min at RT to inactivate SDS. Subsequently, 100 U of HpaII was added and the DNA was digested at 37 °C for 2 hr. The reaction was cooled on ice before labelling cleavage overhangs with Biotin-14-dCTP (ThermoFisher) in a 60 uL reaction (digestion reaction, 0.9 μL dGTP, dTTP and dATP at 2 mM each, 4.5 μL 0.4 mM biotin-14-dCTP, 1.2 μL of Klenow Large fragment [5000 units/ml]) for 45 min at room temperature. Reactions were quenched by adding 0.125% final concentration of SDS and incubated for 5 min at room temperature. Blunted DNAs were ligated in dilute conditions as follows: A master mix containing 75 µL 10% Trition-X 100, 100 µL 10X T4 Ligase Buffer (NEB), 2.5 µL of 20 mg / mL BSA (NEB), and 800 µL water per blunting reaction was assembled and chilled on ice. This chilled ligase mix was added to the blunted DNAs and incubated for 15 min at room temperature to inactivate SDS. Ligation reactions were then chilled on ice for 15 min before addition of 3 µL T4 DNA ligase (NEB, 2, 000, 000 U / mL). Reactions were mixed well and incubated at 10-16°C for 4 hr.

After ligation, 10 mM final concentration EDTA and 2.5 µL of 20 mg / mL Proteinase K (NEB) was added, and reactions incubated at 65 °C overnight to digest proteins and reverse crosslinks. DNA was then extracted twice with phenol:chloroform:isoamyl alcohol, isopropanol precipitated, and then the entirely of the DNAs from the two original aliquots was resuspended in 60 µL of water. Next, non-ligated biotinylated DNAs were eliminated in a 100 µL reaction (60 µL purified DNA, 10 µL Buffer #2 (NEB), 0.5 µL 20 mg / mL BSA (NEB), 5 µL each of 2 mM dATP and dTTP) with 0.5 µL of T4 DNA polymerase for 2 hr at 12 °C. DNA was sheared using a Bioruptor (Diagenode) and size-selected by gel extraction to be between 200 and 500 bp.

Paired-end Illumina libraries were generated as in (Guo et al, 2021) as follows using DNA LoBind tubes (Eppendorf) (13). First, DNA was end-repaired with 4 µL T4 DNA polymerase (NEB), 4 µL T4 PNK (NEB), and 0.75 µL Klenow large fragment (NEB) in 100 µL T4 DNA ligase buffer with 0.25 mM final concentration dNTPs for 30 min at room temperature. Repaired DNA was recovered by Ampure XP (Beckman Coulter) bead purification, using 100 µL beads in 300 µL 20% PEG/NaCl solution. Beads were washed twice with 80% ethanol, dried, and resuspended in 32 µL EB (10 mM Tris-HCl [pH 8.0]). The bead slurry was directly treated with 3 µL Klenow (3’→5’ exo-) (NEB) in 40 µL reaction with 0.2 mM ATP at 37 °C for 30 min to add 3’ overhangs to DNA. Repaired DNA was recovered by Ampure XP capture and resuspended in 15 µL EB without beads. Y-shaped adaptors were prepared by annealing Illumina PE adapter 1 and Illumina PE adapter 2. Y-shaped adapters were added to bead slurry, and the mix was ligated in 25 µL total volume with 5 µL 15 µM ligated adapters and 1.5 µL T4 DNA Ligase (NEB, 400, 000 U / mL) for 30 hr at room temperature, with occasional mixing. Biotin-labeled junctions were purified from non-labelled junctions with Dynabeads MyOne Streptavidin C1 (ThermoFisher). Dynabeads were first washed twice with 1X NTB (5 mM Tris-HCl [pH 8.0], 0.5 mM EDTA and 1 M NaCl) then added to ligation and mixed well. Biotin-labeled DNAs were allowed bind for 30 min at room temperature with occasional mixing before pulldown with a magnet. The beads were washed three times with 100 µL 1X NTB, then washed with 100 µL water, resuspended in 10 µL water, transferred a new tube, and stored at -20 °C. DNA libraries were amplified in 50 µL final volume with 2X KAPA HiFi Master Mix (Roche) supplemented with 5 % final concentration DMSO (Sigma) and appropriate barcoded primers. The total number of cycles was optimized for each sample to minimize the number of cycles required for library generation. Libraries were purified by two-step Ampure XP capture to recover 200-500 bp amplified libraries. DNA was recovered from Ampure beads by resuspending in 20 µL 10 mM Tris-HCl (pH 8.0). Paired-end sequencing was performed with a 91nt kit on a NextSeq500 at the MIT Bio Micro Center.

## RESULTS

### Positive supercoiling sites correlate with higher-than-average Hi-C contact frequencies in *E. coli*

Although supercoiling is known to alter DNA shape, how supercoiling underlies the three-dimensional organization of bacterial chromosomes is unknown. To understand how supercoiling contributes to chromosomal organization, we first sought to compare *E. coli* chromosome architecture deduced from Hi-C mapping with a recently published map of *E. coli* positive supercoiling using GapR-seq (LB, 37°C) (13). From the GapR-seq data, we identified 609 reproducible GapR-binding peaks (see **Material and Methods**, ChIP-seq data processing) along the *E. coli* genome. Because GapR preferentially binds to overtwisted double-stranded DNA (dsDNA), these peak positions correspond to positively supercoiled sites (13, 15) and hereafter we term them as PS (positive supercoiling) sites. Next, we integrated these PS-sites with three chromosome architecture datasets: two publicly available 5C and Hi-C datasets (referred to as Hi-C data below) (4, 5), in addition to one independent Hi-C experiment we performed in this study. We divided the *E. coli* genome of 4.6 million base (Mb) pairs into either 928 (5 Kb per bin) or 9284 (0.5 Kb per bin) bins based on the maximum resolution of the respective Hi-C datasets (**Fig. 1B**). Each 5 Kb bin and 0.5 Kb bin was then assigned a tag, PS/aPS (presence/absence of a PS-site, see **Material and Methods**, HiC/5C bin classification based on ChIP-seq signals and **Supplement Fig. S1D, Table 1** for summary statistics of ChIP-seq peaks).

When we compared the chromosome contacts of bins containing positive supercoiling (PS, **Fig. 1C, D**, dense black bars) to those lacking positive supercoiling (aPS, **Fig. 1C, D**, sparse black bars), we found that PS bins appeared to have high contact frequencies with genomic loci hundreds of kilobases away than aPS bins. To qualitatively illustrate this observation, we calculated the sum of the contacts for each locus within a 100 Kb to 1 Mb window (purple lines in **Fig. 1C, 1D**) and observed that the total contacts are higher in genomic regions with dense PS-sites (at ∼3 Mb in the genome) and lower in the region with sparse PS-sites (at ∼1 Mb in the genome) (**Fig. 1E, 1F)**. We validated our observation in three other 5C/Hi-C experimental replicates at both 5 Kb and 0.5 Kb resolution (**Supplement Fig. S1A**). The length scale (∼ Mb) of the separation between PS and aPS sites matches the scale of the *E. coli* macrodomains (8).

To quantify? the differences in DNA association between positively supercoiled and non-positively supercoiled loci, we computed the inverse of the normalized Hi-C contact frequency between any two bins and plotted the averages against the linear genomic distance between the loci pairs (**Fig. 1G, H**). This inversed Hi-C contact count serves as a proxy for the spatial separation because it is proportional to the spatial distance between two chromosomal sites (27). Our results show that this apparent spatial separation between two PS bins (**Fig. 1G, 1H,** PS-PS, blue) was smaller than that of all bins at all separation ranges (all-all, red). Conversely, the spatial separation between two aPS bins (**Fig. 1G, 1H**, aPS-aPS, green) was larger than the spatial separation of all bins at all distances. The relative difference in the spatial separation between the PS-PS and aPS-aPS curves (([aPS-aPS] - [PS-PS])/max([aPS-aPS] - [PS-PS])) peaks near the 500 Kb loci separation distance, remaining high until 1 Mb, and then gradually decreases (**Fig. 1I, 1J**). We observed a similar trend and pattern on other 5C/Hi-C replicates conducted in different growth conditions and with different resolutions (**Supplement Fig. S1A-B**). Taken together, these data support our observation that positively supercoiled loci interact more frequently than sites that are not positively supercoiled and that this effect is most significant at a macrodomain scale (500 Kb – 1 Mb).

Next, to verify that the observed pattern is indeed dependent on positive supercoiling, we re-analyzed the Hi-C data using a mutant GapR (GapR^1-76^) dataset. GapR^1-76^ lacks the GapR dimerization domain required for tetramerizing on positively supercoiled DNA, and hence binds to DNA nonspecifically (13, 15). GapR^1-76^ binding sites are distributed uniformly along the *E. coli* genome (**Fig. 1C, 1D**, green bars) unlike GapR, which preferentially binds surrounding the *oriC* half of the chromosome (see **Supplemental Table 1A** for definitions of *ori*C half and *ter* half). The difference in spatial separation between GapR^1-76^-bound bins (PS^1-76^-PS^1-76^) and GapR^1-76^ unbound bins (aPS^1-76^-aPS^1-76^) is substantially diminished, validating that PS sites are specific to GapR binding and that they are associated with long-distance contact (**Supplement Fig. S1C**). In summary, our analyses reveal that positively supercoiled loci make more frequent spatial contacts than non-positively supercoiled loci, with the largest effect on spatial association at a macrodomain scale.

### Negative supercoiling sites correlate with higher-than-average Hi-C contact frequencies in *E. coli*

Bacterial genomes are typically net-negatively supercoiled (27), and previous research suggests that this type of supercoiling is a main contributor for genome compaction (28). To explore how negative supercoiling affects chromosome organization, we used an *Ec*TopoI-seq data set in which all *E. coli* TopoI binding sites were identified using immunoprecipitation under the same growth condition as the 5 Kb Hi-C datasets (LB, 37°C) (14). *E. coli* TopoI removes excessive negative supercoiling and was proposed to mark sites of negative supercoiling in *E. coli* (14). Because TopoI can also bind to RNA polymerase during transcription (16), we excluded TopoI binding sites within a transcription unit (TU), in order to focus exclusively on TopoI binding to negative supercoiling. Using the EcTopoI-seq dataset, we identified 446 reproducible TopoI-binding sites and assigned accordingly the 5K *E. coli* genome bins as either NS (negative supercoiling, 38% of total bins) or aNS (absent of a NS-site, 64% of total bins) similarly as that for PS sites (**Fig. 1A, B**, see **Material and Methods**, ChIP-seq data processing & HiC/5C bin classification based on ChIP-seq). NS bins along the chromosome have a largely similar distribution as PS bins in that they are densely distributed surrounding the *ori*C half of the chromosome (**Supplemental Table 1A**, *ori*C half and *ter* half), which contains multiple *rrn* operons, and sparsely surrounding the *ter* half of the chromosome. Similar to what we observed for PS bins, genomic regions with more frequent NS bins (**Fig. 1C, 1D,** dense blue bars) also showed a higher Hi-C contact frequencies with other genomic loci a few hundred Kb away than the ones with sparse NS bins (sparse blue bars).

To quantify differences in DNA association between negatively supercoiled loci (NS) and those absent of negatively supercoiled sites (aNS), we calculated the apparent spatial separation between two negatively supercoiled bins along the chromosome (**Fig. 2A-2B**). Akin to our previous observations with positive supercoiling, we found that the spatial separation between two bins containing negatively supercoiled DNA (NS-NS, blue) is the lowest. This is followed by the spatial separation between all genomic locations (all-all, red), with the largest spatial separation occurring between two non-negatively supercoiled bins (aNS-aNS, green). Furthermore, the relative difference in spatial separation between NS-NS and aNS -aNS sites follows the same trend as observed in the GapR analysis (**Fig 2C, D**). These observations are also reproducible across all Hi-C experimental replicates, despite differences in the data resolution, experimental conditions, and technologies. (**Supplement Fig. S2A-B**). Our analyses here suggest that, as with positive supercoiling, two negatively supercoiled loci make more frequent spatial contacts than non-negatively supercoiled loci, again with the largest effects on spatial association at macrodomain scales.

**Figure 2.**
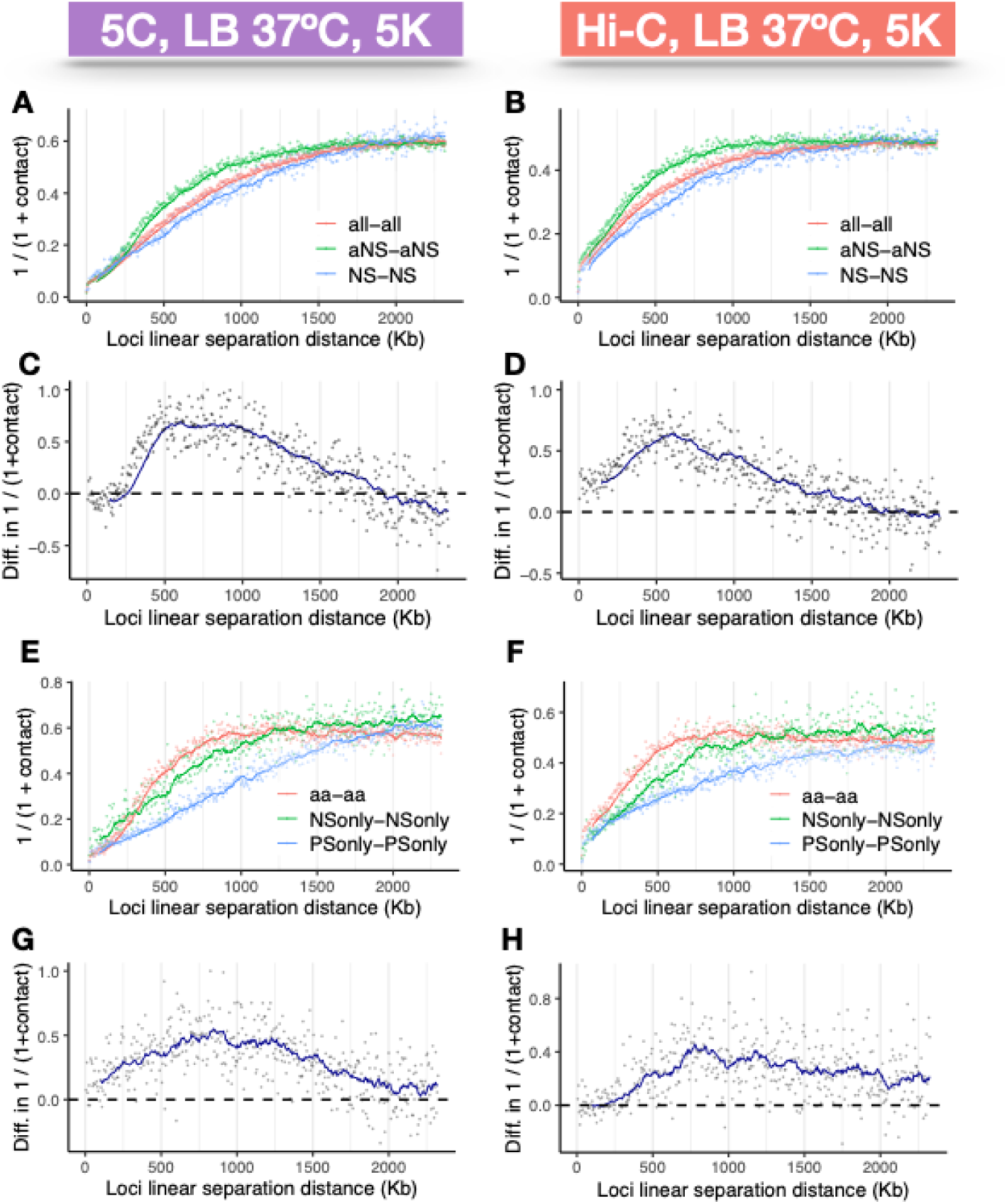
*E. coli* chromosomal loci containing negative supercoiling sites exhibit higher-than-average but lower spatial contacts in Hi-C maps than loci containing positive supercoiling. **(A-B)** Inversed average contact frequencies of two genomic loci (*y*-axis) at each linear genomic separation distance (*x*-axis) with simple moving averages (colored curves, window size = 15) for loci pairs both containing negative supercoiling sites (NS-NS, blue), not containing positive supercoiling sites (aNS-aNS, green), and all loci pairs (all-all, red) show that NS-NS loci pairs have on average lowest inversed average contact frequencies (highest spatial contact) at all linear genomic distances than aNS-aNS or all-all loci pairs. **(C-D)** Numeric difference in the inversed average contact frequencies (y-axis) between NS-NS loci and aNS-aNS loci at each genomic separation distance (x-axis) reaches maximum in the range of 500-1000 Kbp Kbp. **(E-F)** Inversed average contact frequencies of two genomic loci (*y*-axis) at each linear genomic separation distance (*x*-axis) with simple moving averages (colored curves, window size = 15) for loci pairs containing positive supercoiling site only (PS only -PS only, blue), negative supercoiling site only (NS only – NS only, green), and none (aa – aa, red) show that PS only-PS only loci pairs have on average lowest inversed average contact frequencies **(G-H)** Numeric difference in the inversed average contact frequencies (y-axis) between PS only-PS only loci and aa-aa loci at each genomic separation distance (x-axis) reaches maximum in the range of 500-1000 Kbp.

### Positive supercoiling has a larger effect on chromosomal DNA interaction than negative supercoiling

Our analyses so far suggest that both positive and negative supercoiling play significant roles in *E. coli* chromosome compaction (**Fig. 1G, H, 2C, D**). Interestingly, when we compared the apparent spatial separations (for example, compare the y axes of **Fig. 1G** and **Fig. 2A**), positive supercoiling appeared to have a greater effect than negative supercoiling, i.e. PS-PS spatial separations appear to be closer than those of NS-NS sites. To quantify this difference, we assigned each 5 Kb or 0.5 Kb genomic bin a label depending on what type of supercoiling, if any, dominants within that bin (PS-only, NS-only), or is absent of either (aa) (see **Material and Methods**, HiC/5C bin classification based on ChIP-seq signals). Bins containing both GapR and TopoI peaks were not used in the analysis (a total of 162 out of 928 5 Kb bins and 539 out of 9284 0.5 Kb bins). This mutually exclusive bin assignment allows us to isolate the effects of each supercoiling type in our analysis. Plotting the apparent spatial separation within these three groups confirmed that positively supercoiled bins (**Fig. 2E, F**, PSonly-PSonly, blue) associate more frequently with each other than negatively supercoiled bins (NSonly-NSonly, green), with non-supercoiled loci displaying the lowest frequency of association (aa-aa, red). Interestingly, at very long distances (>1.25 Mb), the spatial separation of non-supercoiled bins (aa-aa) was slightly smaller than negatively supercoiled bins and approached or became even smaller than positively supercoiled bins at the largest linear distances (> 2 Mb). The significance of these differences is unclear and may require further investigation. Importantly, the spatial separation between two positively supercoiled loci is always lower than that of two negatively supercoiled loci, as demonstrated by the positive difference between PSonly and NSonly’s apparent spatial separation (**Fig. 2G, 2H**). Similar patterns were found in all other independent Hi-C replicates (**Supplement Fig. S2B**). Based on these analyses, we conclude that while positive and negative supercoils both package the chromosome, positive supercoiling leads to higher spatial contacts than negative supercoiling, especially for contacts that occur on the scale of macrodomains (∼500 Kb to ∼1.5 Mb, **Fig. 2G, 2H**).

### Positive, but not negative, supercoiling sites can be inferred from Hi-C contact map

Our results suggest that positive and negative supercoiling generate distinct chromosome organizational patterns that are captured in the Hi-C map. We wondered if supercoiling is a strong enough underlying feature of chromosome organization that the locations of supercoils can be inferred from Hi-C data. To investigate whether the Hi-C contact matrix encodes such supercoiling information, we utilized a random forest model to predict positively and negative supercoiling loci based on the corresponding contact frequencies. We chose random forest model because its prediction performance is robust under limited sample points and a large number of covariates.

Our random forest models were first trained using the transformed Hi-C contact matrices and the correct supercoiling label of each genomic loci (bin size = 5 Kb) obtained from the ChIP-seq data sets (see **Material and Methods**, Training the random forest models and **Fig. 3A**, left). In the training, each row of the input feature is the contact frequencies of a genomic locus with itself (brown square) and all other flanking loci on both sides (blue squares); the output, or response, is whether this locus contains a PS peak, NS peak, both, or none (green and purple boxes on the right). After the models were trained with varying amounts of data, we ask the models to predict PS/aPS and NS/aNS bins for a set of unseen, randomly distributed bins, based solely on their Hi-C contact frequencies with all other genomic loci (**Fig. 3A**, right). To evaluate the model prediction performance, we compared the average AUROC and AUPRC computed on the testing set with those of the baseline models, which were trained on randomly assigned supercoiling labels. The ROC (Receiver Operating Characteristic) curve plots the true positive rates against the false positive rates at different decision thresholds for making a prediction, and AUROC is the area under the ROC curve, which is commonly used to evaluate the performance of a prediction model. An AUROC value of 1 means a perfect prediction model, and values between 0.5 and 1 indicates the prediction is better than a random prediction model. The precision-recall curve (PRC) plots the positive predictive values (precision) against the true positive rates (recall) for different threshold values, and AUPRC summarizes the area under the precision-recall curve, which is an alternate metric for evaluating the prediction performance. The AUPRC ranges between 0 and 1, where higher values indicate better model performance. A value of 0 indicates that the model’s precision is always zero, regardless of the recall, and a value of 1 indicates perfect precision-recall trade-off. (see **Material and Methods**, Evaluating random forest model prediction performance). The combination of the two parameters allows the model classification to be robust against potential class imbalance issues, for example, the numbers of NS and aNS bins are not equal.

**Figure 3.**
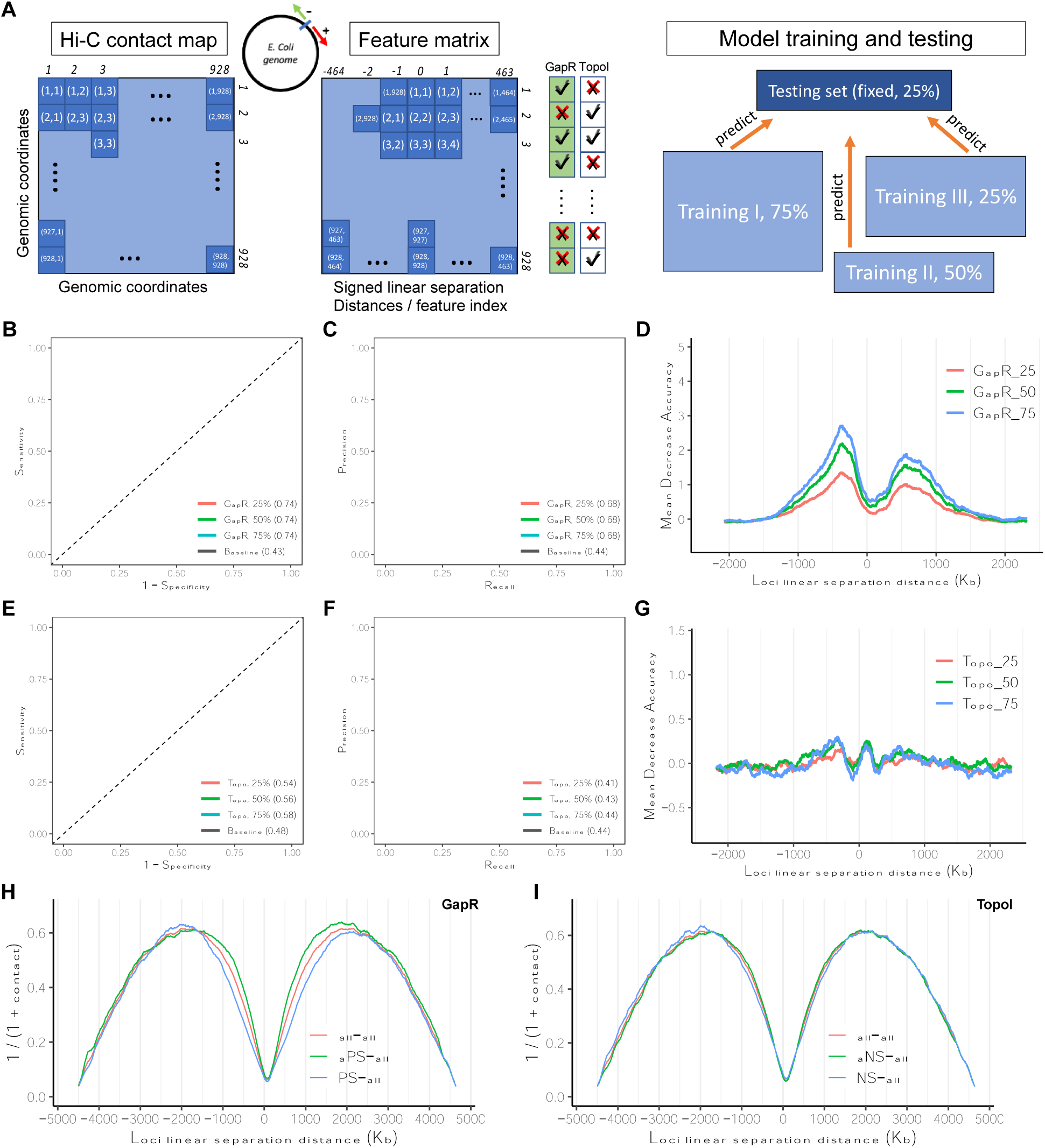
Machine learning models can predicate positive but not negative supercoiling sites accurately from Hi-C contact map. (**A**) A cartoon illustration of the machine learning model design. The input matrix is a rearrangement of the conventional contact matrix^7^ (left), in which each input feature (row) is the contact frequency of a genomic locus (one 5 Kbp bin, brown) with flanking chromosome on both sides (-464 to + 463 bins, blue). The responses, or whether this 5 Kb bin contains any Gap (green) or TopoI (purple) peaks, are indicated by the check and error marks (middle). A set of randomly chosen loci (25% out of 928 bins) is preserved as a fixed testing set for all random forests models (right). (**B-C**) ROC and PR curves of 10 independent experiments (light-shaded curves) with 25% (red), 50% (green), and 75% (blue) training data predicted positive supercoiling (marked by GapR peaks) locations robustly with similar AUROC (0.74) and AUPRC (0.68) values, while the baseline models (black), trained with randomized peak labels, are at ∼ 0.43 and 0.44 respectively. (**D**) Average feature importance (over 10 independent experiments) measured in mean decrease accuracy (colored dots, with simple moving averages of window size = 15 as solid curves) from experiments with 25% (red), 50% (green), and 75% (blue) training data peaked at ∼ 500 Kb genomic separation distance, indicating that these features contribute the most to the effective classification of GapR peak-containing bins. (**E-F**) ROC and PR curves of 10 independent experiments (light-shaded curves) with 25% (red), 50% (green), and 75% (blue) training data did not predicate negative supercoiling (marked by Topo I peaks) locations robustly. The AUROC (0.54-0.58) and AUPRC (0.41-0.44) values are not significantly different from those of the baseline models (black curves), indicating that the models cannot effectively recover negative supercoiling spatial patterns from Hi-C contact maps. (**G**) Average feature importance (over 10 independent experiments) measured in mean decrease accuracy (colored dots, with simple moving averages of window size = 15 as solid curves) from experiments with 25% (red), 50% (green), and 75% (blue) training data showed uniform patterns across different genomic separation distances (**H**) The average contact frequencies (y-axis, inversed contact counts representing the apparent spatial separation between two loci) at each signed genomic separation distance (x-axis) are plotted (colored dots) with simple moving averages (colored curves, window size = 15). Different colors indicate the interaction between loci pairs with at least one containing positive supercoiling site (PS-all, blue) and that with at least one containing no positive supercoiling site (aPS-all, green). The average contact frequencies at each genomic separation are plotted in red as a reference. (**H**) The inverse average contact frequencies among PS-all (blue), aPS-all (green), and all-all (red) loci pairs in the range of ± 500 – 1000 Kbp are significantly different from each other, explaining the effective prediction of positive supercoiling sites on *E. coli* genome using Hi-C contacts as the input. (**I**) The inverse average contact frequencies among NS-all (blue), aNS-all (green), and all-all (red) loci pairs in all ranges are not significantly different from each other, explaining the ineffective prediction of negative supercooling sites on *E. coli* genome using Hi-C contacts as the input.

We observed that the random forest model performed well for predicting positive supercoiling using 75% of data for training, achieving an AUROC value of ∼0.74. The model’s prediction performance remains consistently robust (AUROC = ∼0.74) even when less data was used for training (50% and 25%), outperforming the baseline models (where the target labels were permuted) where a low AUROC value of 0.43 was observed (**Fig. 3B**, and see baseline models definition **in Material and Methods,** Evaluating random forest model prediction performance, Training the random forest models). Using AUPRC as the evaluation metric yielded similar results, where our models trained with different data amounts achieved an AUPRC of ∼0.68, as compared to AUPRC of 0.44 for the baseline models (**Fig. 3C**). These consistently robust performances across models with different training-to-testing data ratios suggest that positively supercoiled loci have highly uniform, strong interaction patterns with other genomic loci, and that such patterns distinguish them from loci without positive supercoiling.

To gain a deeper understanding of how the models can distinguish between PS (containing positive supercoiling sites) and aPS (absent of positive supercoiling sites) bins, we computed the feature importance using the Mean Decrease in Accuracy (MDA), a metric that measures how much accuracy the model losses by excluding each feature. Features with high MDA values (in this case, Hi-C contact frequencies at a certain linear distance along the chromosome) significantly contributes to the classification of the associated genomic locus to PS or aPS. The MDA plot peaked at ∼500 Kb, indicating that spatial contact between loci separated by this distance is the greatest contributor to the correct categorization of PS and aPS bins (**Fig. 3D**). This result confirms our finding that positively supercoiled and non-positively supercoiled bins differ the most in their interactions with loci ∼500 Kb away, and the spatial interaction patterns at these separation ranges can robustly predict the existence of positive supercoiling at any genomic loci.

Next, to determine whether Hi-C data can be used to predict negative supercoiling in a similar fashion, we trained the random forest models using TopoI-seq data, following the same procedure as described above. Surprisingly, we found that the model’s performance was not as good as predicting positive supercoiling, with AUROC ranging from 0.54 to 0.58 at different training-to-testing data ratios. AUPRC scores from these models ranged from 0.41 to 0.44, which are indistinguishable from or slightly worse than the baseline model with AURPC at 0.44 (**Fig. 3E, 3F**). Furthermore, the MDA plot did not show a clear pattern as the positive supercoiling prediction model (**Fig. 3G**). These results indicate that Hi-C contact frequencies contain some information but cannot effectively determine whether a locus is negatively supercoiled, in contrast to what we observed with positive supercoiling (**Fig. 3B-3D**). Together, these observations indicate that positive supercoiling enhances chromosomal loci spatial interactions more than negative supercoiling.

We conclude this section by examining the reasons that explain the different prediction performance of PS and NS peaks. The presence of significant difference between the interactions of PS and aPS loci with all genomic loci (**Fig. 3H**, green and blue lines, **Supplement Fig. S3A**) explained the robust prediction performance of PS sites. The poor performance of TopoI peak prediction could be attributed to the lack of significant difference between the interaction pattern of NS and aNS with all genomic loci (**Fig. 3I**, overlapping green and blue lines, **Supplement Fig. S3B**). Although there is a significant difference between NS-NS and aNS-aNS interactions (**Supplement Fig. S3B**, green and purple lines), this NS-all versus aNS-all interaction difference was not different from each other accessible to the models as the features were not labeled as peak or non-peak. Instead, the models rely on the difference between peak and non-peak with all genomic loci to make a classification (**Supplement Fig. S3B**, yellow and blue lines). Since such a difference (peak-to-all versus non-peak-to-all) is significant in PS sites but not NS sites, we conclude that recovering positive supercoiling signals, but not negative supercoiling, is possible from using Hi-C datasets alone. Our findings suggest that positive supercoiling has a characteristic signature that is reflected in the Hi-C contact map.

### Transcription-independent and -dependent supercoiling

Transcription is a major supercoil-generator in bacteria (29, 30). During transcription, an elongating RNA polymerase (RNAP) under-twists upstream DNA and over-twists downstream DNA, generating negative and positive supercoiling, respectively (10). By examining the positions of genomic supercoiling relative to the transcriptional activity of nearby transcription units (TUs), we can access the capacity of transcription to generate persistent supercoiling and consequently its contribution to genome packaging. We used a set of RNA-seq data (LB, 37°C) with 2640 TUs annotated on the *E. coli* chromosome. We calculated the distance between a transcription end site (TES) and the nearest PS site indicated by a GapR peak downstream, and the distance between a transcription start site (TSS) and its nearest NS site (TopoI peak) upstream. The distributions of the distance between NS/PS site to its nearest TSS/TES do not differ from each other significantly (p=0.13, Wilcoxon, **Fig. 4A, B**). The majority (94.7% and 94.8%) of PS and NS sites are within 1.0 Kb of its nearest TES and TSS, respectively. For comparison, on average 90.0% and 77.0% of random chromosomal sites are within the same 1.0 Kb distance from the nearest TES or TSS, respectively, due to the close spacing between Tus along the *E. coli* chromosome (**Fig. 4C, D** & **Supplement Fig. S4B, S4C**). This comparison demonstrates that the majority of supercoils are associated with transcription activities, and that it makes up to ∼95% of total sustained genomic supercoils, hence transcription contributes significantly to genome organization.

**Figure 4.**
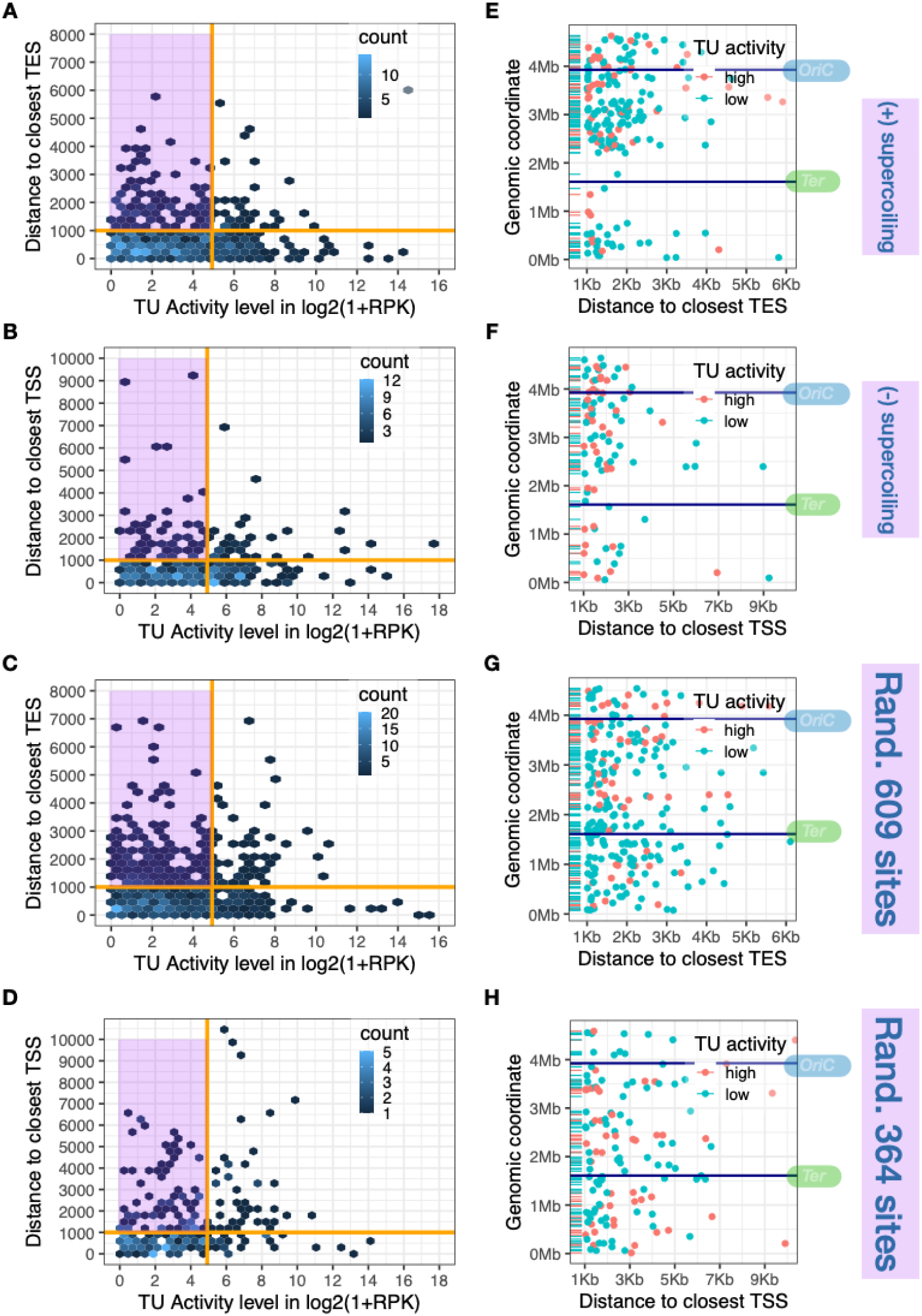
Transcription is a strong driving force for supercoiling generation. (**A**) 2D-histrogram showing the distribution of supercoiling sites (hexagon bins colored by density) with respect to the distance (bp, y-axis) to their nearest upstream transcription end site (TES) and the normalized (log2) transcription activity (x-axis, RPK) of that transcription unit (TU). Orange vertical line indicates the 75^th^ percentile of all TU activities (x-intercept = 29.34). Orange horizontal line indicates 1 Kb away from the nearest TES (y-intercept = 1000 bp). Positive supercoiling sites located in the purple block are 1 Kb away from the nearest TES and the associated TUs have transcription activities below the 75^th^ percentile, suggesting that those sites are not associated with transcription. (**B**) The negative supercoiling counterpart of (**A**). (**C-D**) Control experiments repeating the analysis in (**A and B**) by randomly generating a set of genomic locations matching the number of PS and NS sites. More randomized sites reside within the purple blocks (> 1 Kb away from TUs of low transcription activities) than those in (**A-B**). (**E**) PS sites (colored dots) greater than 1 Kb away from their nearest TES (x-axis) are plotted to visualize their location along the *E. coli* chromosome (y-axis). Position of *Ori*C and *Ter*C are indicated by two horizontal lines. Positive supercoiling sites are colored in red if they associate with a highly transcribed TU (top 25% percentile) and in green otherwise. (**F**) The NS counterpart of (**E**). (**G-H**) Control experiments repeating the analysis in (**E**) using the randomly generated sets of genomic sites.

Because all TUs make up to 92% of the *E. coli* genome in length (31), ∼ 45% of the TU coverage overlap with either PS or NS sites (identified by GapR and TopoI ChIP-seq binding sites, respectively), but 65% of these overlapping regions are associated only with PS sites. Previous studies have shown that DNA supercoiling generated by active transcription may modulate the activity of RNA polymerase molecules on the same gene as well as impact the transcription of adjacent genes (31, 32, 33). Therefore, the observation that positive supercoils extend into adjacent TUs more frequently than negative supercoils suggests that positive supercoiling may play a larger role than negative supercoiling in regulating *E. coli* transcription. As transcription is largely suppressed by positive supercoiling but promoted by negative supercoiling, this differential regulation could maintain most TUs in a repressed state, which only turns on when needed to minimizing potential transcription cost.

We also observed that a minor subset of GapR and TopoI peaks (3.8% and 4.1% of all peaks, respectively) are located more than 1 Kb from any TUs (indicated by hex bins above the orange line, **Fig. 4A, 4B**). Approximately 72% and 79% of this subset associate with TUs with low transcriptional activities, respectively (expression level below the 75^th^ percentile of all TUs, purple-highlighted quadrant, **Fig. 4A, 4B**). Furthermore, we observed that this minor group tend to be located near the *Ori* and *Ter* regions but are sparse in between (**Fig. 4E, 4F**). These patterns were not observed in the control analysis of randomly selected chromosomal sites, which were uniformly distributed across the genome as expected (**Fig. 4G, 4H**). These observations suggest that these supercoils may be introduced by genomic processes other than transcription since they are far from highly active TUs (expression above the 25^th^ percentile of all TUs), but likely by replication and/or the actions of NAPs and topoisomerases (34).

Taken together, our analyses show that supercoils are mostly associated with transcription activities, in line with previous findings and suggest that transcription is a main driving force for genome structuring.

## DISCUSSION

In this work, we integrated two recently published maps of *E. coli* positive and negative supercoiling (GapR-seq and TopoI-seq, respectively) with three independent Hi-C data sets to explore computationally the roles of positive and negative supercoiling in *E. coli* genome organization (**Fig. 1C-J**). We observed a consistent, robust pattern across various Hi-C data sets that supercoiled genomic loci have shorter spatial separation than non-supercoiled sites, with two positively supercoiled loci having shorter spatial separations than two negatively supercoiled loci (**Fig. 2A-D, 2E-H**). Both positive and negative supercoiling have the largest effects on spatial association of loci are separated by 0.5-1.5 Mb in the linear chromosome, on the order of the *E. coli* macrodomains. Building upon these observations, we developed a supervised machine learning model and found that Hi-C data can accurately predict positively supercoiled sites but not negatively supercoiled sites, suggesting that the spatial interactions mediated by positive supercoiling dominate Hi-C measurement (**Fig. 3A-I**). Finally, we found ∼ 95% of GapR and TopoI sites reside next to a TU, and that more positive supercoiling sites than negative supercoiling sites are within the body of an adjacent TU. These observations suggest that transcription is the major driver of supercoiling and genome organization. Taken together, our results support the longstanding idea that supercoiling leads to chromosome packaging and suggest that, although positive and negative supercoiling can be reductively considered to structure DNA similarly, their effects on chromosome organization are not equivalent. Our work suggests that positive supercoiling enhances spatial contact more than negative supercoiling, which has not been proposed (**Fig. 4A-F**).

### The differential roles of positive and negative supercoiling in chromosome organization

The *E. coli* chromosome is naturally negatively supercoiled (27), which is necessary to allow genomic processes such as replication or transcription to proceed without being energetically costly to open the double-stranded chromosomal DNA. As such, it is reasonable to expect that positive supercoiling is necessary to package the genome and compensate naturally existing negative supercoiling, which may destabilize the chromosome by randomly introducing ssDNA regions due to stochastic dsDNA melting.

Why would positive and negative supercoiling have differential effects on chromosome organization? The mechanisms that regulate positive and negative supercoiling in *E. coli* are complex. However, we note that there are many more abundant DNA structuring proteins in *E. coli* (e.g., HU, H-NS, Fis) that are known to constrain negative supercoiling than those constrain positive supercoiling, suggesting that the effects of negative supercoiling may be buffered by these DNA binding proteins (35, 36, 37). Hence, the role of negative supercoiling may be more pertinent to genomic processes while that of positive supercoiling may be more pertinent to genome organization. Further investigation will be needed to address this question.

### Genomic locations of positive and negative supercoiling

When we arranged PS and NS sites along the *E. coli* chromosome, we found that they share similar patterns. Both PS and NS sites are enriched in *Ori* and *ter* regions, but less in between (**Fig. 1C, 1D**). As the *Ori* macrodomain contains most of the active genes, and both *Ori* and *Teri* are involved in DNA replication (8, 28), the dense location of PS and NS sites in these two regions is consistent with the expectation that transcription and replication are major supercoil generators. Between the two, transcription appears to have a more significant role in generating supercoils, as over 95% of GapR and TopoI peaks are associated with TUs. Of note, there are more GapR peaks (609) compared to TopoI peaks (446) (see **Supplementary Table 1** for statistics), which could be due to different capturing efficiency of the corresponding antibody in the ChIP-Seq experiments.

### Short-distance and long-distance compaction

We show that the spatial distance separation between two bins containing PS or NS sites is smaller than those without these sites, with the largest effects on compaction occurring for loci 0.5-1 Mb apart in a linear chromosome (**Fig. 1I-J, 2C-D**). We hypothesize that the fact that supercoiling-dependent compaction peaks at 0.5-1 Mb is a result of the chromosome conformation and is dictated by the macrodomain organization of the *E. coli* genome. Short-distance interactions such as TADs occur on only ∼100 Kb scales (8), which are too short to explain our observations. In contrast, the observed supercoiling-dependent compaction matches the size of macrodomains, which are estimated to be ∼1 Mb (8). While the actual peak is slightly different for each experiment, they all fall in the 0.5-1 Mb genomic distance range. Such a difference may be due to experimental variations since it has an association with the number of valid paired reads from the Hi-C/5C experiment. As such, we hypothesize that positive and negative supercoiling contributes to organizing the chromosome by enhancing the interaction within a macrodomain.

### Physical models of the supercoiling-organized *E. coli* chromosome

What would be the physical model of the *E. coli* chromosome architecture that explains our observations? There are two challenges to forming a unified model. First, most models of chromosome architecture are based on the underlying assumption that physical distance and contact frequency are inversely correlated (7, 38, 39). However, this relationship is not necessarily valid for all loci and, in experiments that test this relationship, the variation between contact frequency and physical distance can be large (7, 39, 40, 41). Second, Hi-C captures population-averaged contact, meaning that contact could result from distant loci being physically closer but static, or from dynamic (and potentially rare), frequent collisions between loci. Therefore, we propose two models, one static and one dynamic. In the static model (**Fig. 5A**), both positive and negative supercoiling lead to the compaction of the chromosome but positive supercoiling has a stronger effect than negative supercoiling. As such, chromosomal sites containing positive supercoils may form the core of the condensed chromosome, while sites containing negative supercoils are on the periphery of the core, allowing genomic process such as replication and transcription to occur. In this sense, positively supercoiled and negatively supercoiled chromosomal sites may be analogous to heterochromatin and euchromatin in eukaryotes. In the dynamic model (**Fig. 5B**), the chromosome may be an amorphous network of DNA that is being dynamically reshaped due to the transitory, nonspecific interactions with nucleoid-associated proteins such as HU (42). Sites containing positive supercoils may be more mobile due to the lack of NAP binding and hence have a higher chance to interact with each other over long linear distances in the nucleoid (**Fig. 5B**, blurry lines). Sites containing negative supercoils and no supercoils, however, are relatively immobile due to the stable association with NAPS, some of them having a higher binding tendency to negatively supercoiled DNA (14), and the transcription/translation machineries, and hence have less chances to interact with each other (**Fig. 5B**, sharp lines and small blobs). Both models should give rise to similar Hi-C contact maps. Further investigation to visualize the spatial localization and dynamics of positively and negatively supercoiled sites in live *E. coli* cells is required to differentiate between these models.

## DATA AVAILABILITY

The sequencing data from this study have been deposited at the Gene Expression Omnibus (GEO): GSE214856. Scripts used for data processing, analysis, trained machine learning models are available from the Xiao Laboratory GitHub repository (XiaoLab JHU: https://github.com/XiaoLabJHU).

## SUPPLEMENTARY DATA

Supplementary Data are available at NAR online.

## ACKNOWLEDGEMENTS

The authors thank all members of the Xiao, Ji laboratories for their valuable discussions and advice on this project. All computations were performed on the Joint High Performance Computing Exchange (JHPCE), organized by the Department of Biostatistics at the Johns Hopkins Bloomberg School of Public Health, and the Maryland Advanced Research Computing Center (MARCC) Rockfish cluster. Z.F. wants to thank Jiafang Song for creating the illustration cartoons and Dr. Hongkai Ji for offering statistical insights.

## SUPPLEMENTAL FIGURES AND TABLES

**Figure S1.**
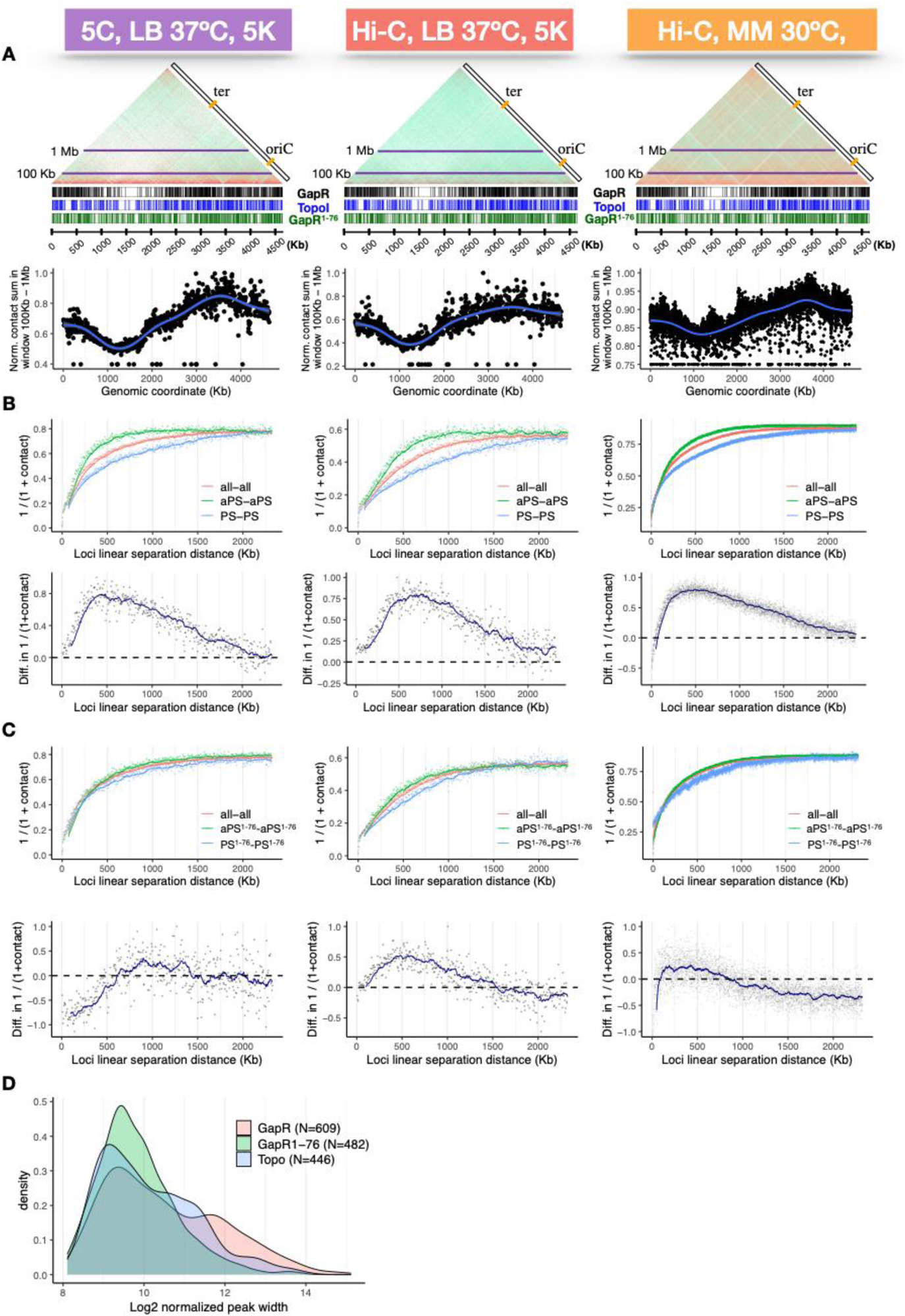
(A) Replicates of the corresponding Hi-C and 5C experiments illustrated in (Fig. 1C-D, E-F) verified the reproducibility of the patterns across different experimental conditions and resolutions as described in the article. (B) Replicates of the corresponding Hi-C and 5C experiments illustrated in (Fig. 1G-H, I-J) verified the reproducibility of the patterns described in the article. (C) GapR^1-76^ counterpart of the analysis illustrated in (Fig. 1G-H, I-J). (D) Density plots for lengths (log2 normalized, x-axis) of identified reproducible GapR peaks (salmon, N=609), GapR^1-76^ peaks (green, N=482), and TopoI peaks (blue, N=446).

**Table S1.** **1A.** The *E. coli* genome is divided into 928 5Kb bins and 4642 0.5 Kb bins. We identified 609 positive supercoiling sites (marked by reproducible GapR peaks) and 446 negative supercoiling sites (marked by reproducible TopoI peaks) (see methods, ChIP-seq analysis). The *OriC* half is defined as the half of the *E. coli* genome centering at the 5 Kb or 0.5 Kb bin which contains *OriC* (coordinate = 3, 925, 744, EcoCyc). The *ter* half is defined by the set of bins complement to the *OriC* half. **1B: Statistics of three sets of identified, reproducible peak widths**

|  | GapR | Topo |
| --- | --- | --- |
| **Total peak number** | 609 | 446 |
| **5Kb bins (928)** | 454 | 352 |
| **0.5Kb bins (9284)** | 3672 | 1890 |
| **5Kb bins in *ter* half (464)** | 147 | 150 |
| **5Kb bins in *OriC* half (464)** | 307 | 201 |
| **0.5Kb bins in *ter* half (4642)** | 714 | 716 |
| **0.5Kb bins in *OriC* half (4642)** | 2958 | 1174 |

**1B: Statistics of three sets of identified, reproducible peak widths**
|  | Min | 1<sup>st</sup> Qu. | Median | Mean | 3<sup>rd</sup> Qu. | Max. |
| --- | --- | --- | --- | --- | --- | --- |
| **GapR** | 278 | 654 | 1193 | 2532 | 3112 | 35457 |
| **TopoI** | 330 | 558.2 | 946.5 | 1647.2 | 1942 | 15616 |
| **GapR<sup>1-76</sup>** | 257 | 583.8 | 820 | 1201.8 | 1302.5 | 14331 |

**Figure S2.**
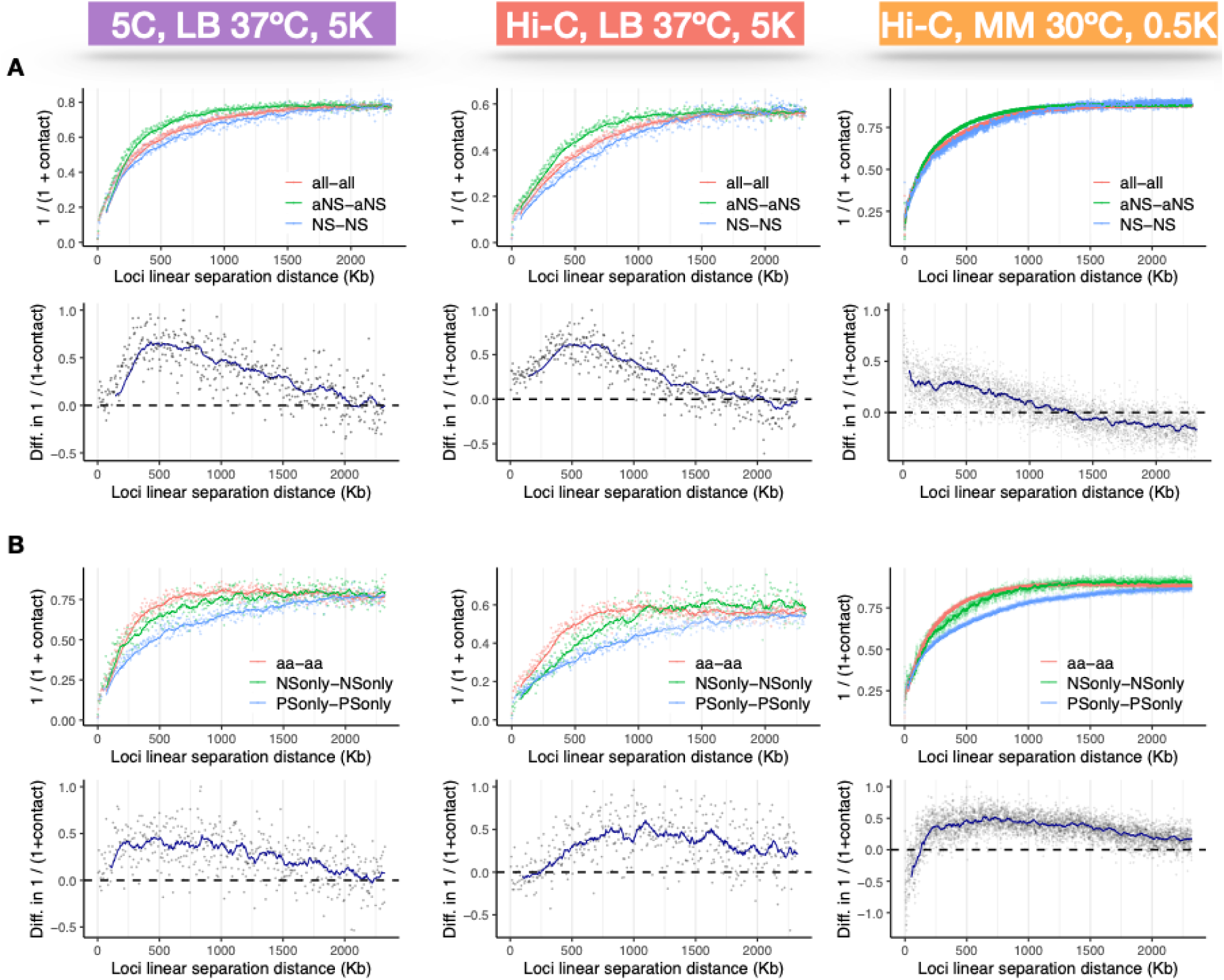
**(A)** Replicates of the corresponding Hi-C and 5C experiments illustrated in (**Fig. 2A-B, C-D**) verified the reproducibility of the patterns described in the article. **(B)** Replicates of the corresponding Hi-C and 5C experiments illustrated in (**Fig. 2F-G, H-I**) verified the reproducibility of the patterns described in the article

**Figure S3.**
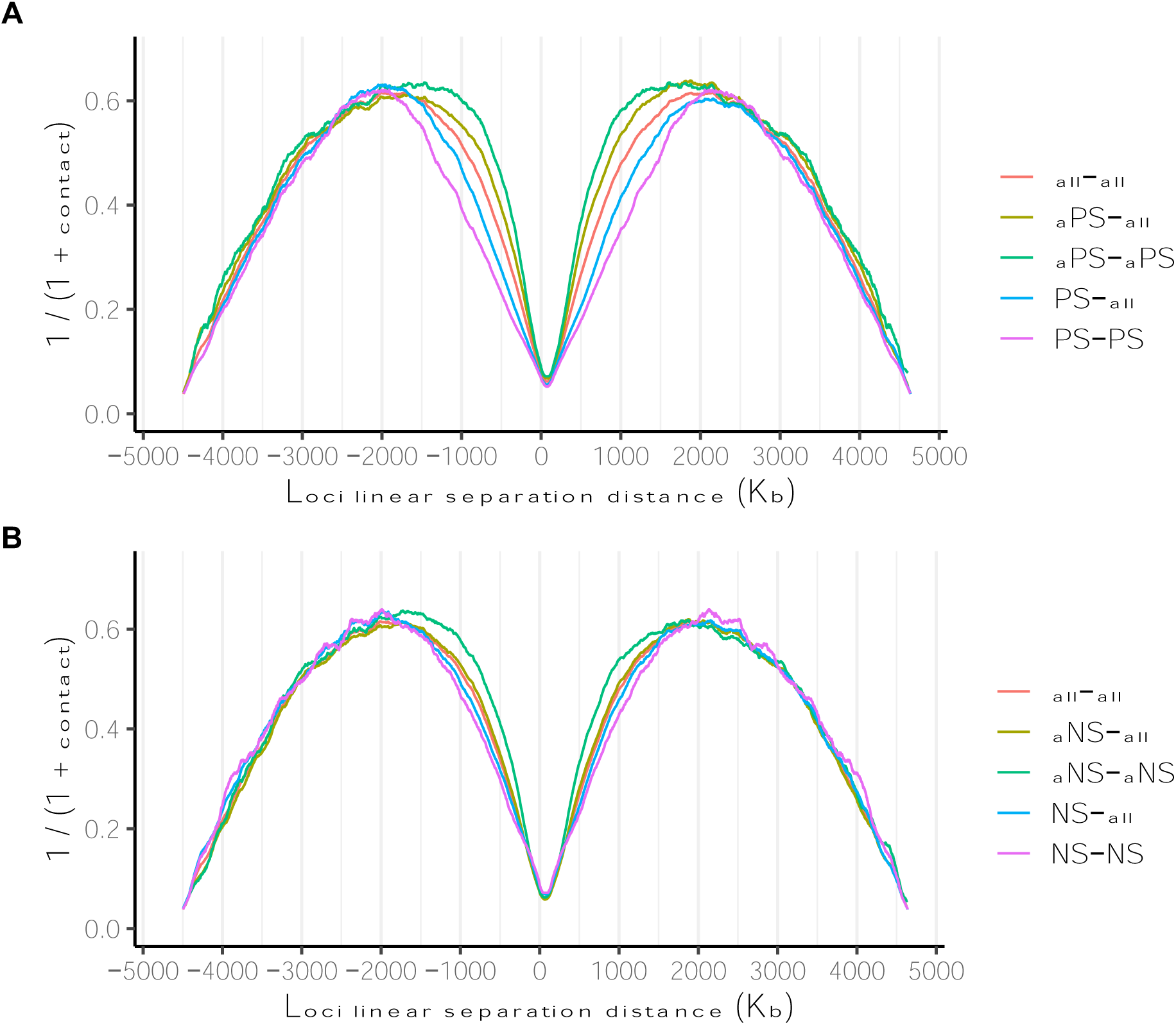
**(A)** Average contact frequencies (y-axis, inversed contact counts representing the apparent spatial separation between two loci) at each signed genomic separation distances (x-axis) are plotted (colored dots) with moving averages (colored curves, window size = 15). Different colors indicate the interaction between all loci pairs (all-all, red), when at least one locus does not contain positive supercoiling site (aPS-all, yellow), when both loci do not contain positive supercoiling site (aPS-aPS, green), when at least one locus contains positive supercoiling site (PS-all, blue), or when both loci contain positive supercoiling site (PS-PS, pink). **(B)** The negative supercoiling counterpart of (A).

**Figure S4.**
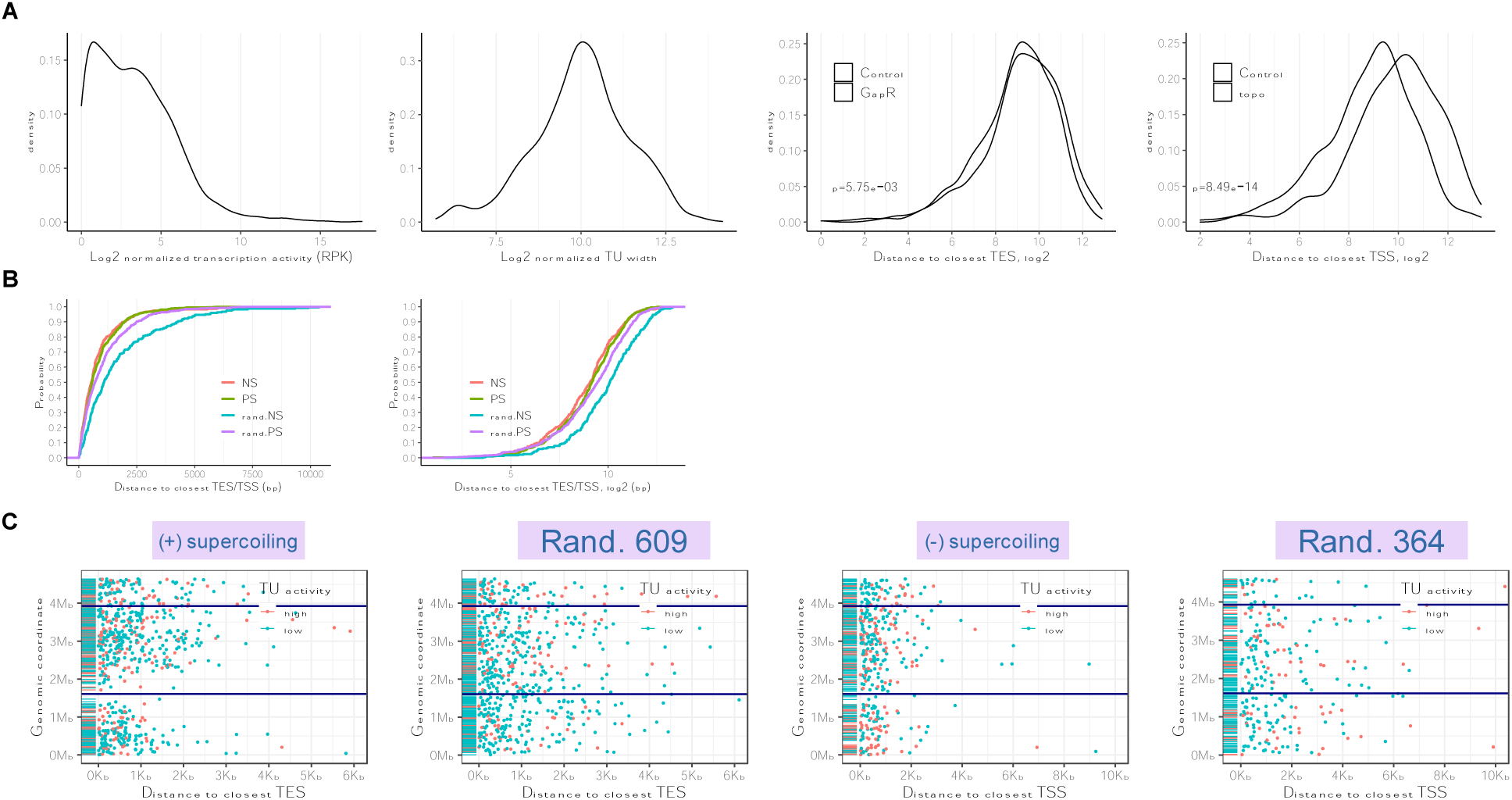
**(A)** Density plots of the log2 normalized transcription unit length (left) and activity (right 2, measured in Reads Per Kilobase). **(B)** Density plots of the distance from PS v.s. random-PS (609 sites) to their nearest TES (top left) and from NS v.s. random-NS (364 sites) to their nearest TSS (top right). The cumulative distribution functions (bottom left) of the four distributions in the above two figures and with log2 normalization (bottom right). **(C)** Four sets of peaks are plotted to visualize their location along the *E. coli* genome (y-axis) and distance to their nearest TU (x-axis). Two horizontal lines in blue indicate the position of *ori*C and *ter*. Positive supercoiling sites are colored in red if they associate with a highly transcribed TU and in green otherwise.

